# Bariatric surgery reveals a gut-restricted TGR5 agonist that exhibits anti-diabetic effects

**DOI:** 10.1101/2020.01.10.902346

**Authors:** Snehal N. Chaudhari, David A. Harris, Hassan Aliakbarian, Matthew T. Henke, Renuka Subramaniam, Ashley H. Vernon, Ali Tavakkoli, Eric G. Sheu, A. Sloan Devlin

## Abstract

Bariatric surgery, the most effective treatment for obesity and type 2 diabetes, is consistently associated with increased levels of the incretin hormone GLP-1 and changes in overall levels of circulating bile acids. The levels of individual bile acids in the GI tract following surgery, however, have remained largely unstudied. Using UPLC-MS-based quantification, we observed an increase in an endogenous bile acid, cholic acid-7-sulfate (CA7S), in the GI tract of both mice and humans after sleeve gastrectomy. We show that CA7S is a TGR5 agonist that induces GLP-1 secretion in vitro and in vivo and that CA7S administration increases glucose tolerance in insulin-resistant mice in a GLP-1 receptor-dependent manner. CA7S remains gut-restricted, minimizing off-target effects previously observed for TGR5 agonists absorbed into circulation. By studying changes in individual metabolites following surgery, this study has revealed a naturally occurring TGR5 agonist that exerts systemic glucoregulatory effects while remaining confined to the gut.

## Introduction

Obesity and type 2 diabetes (T2D) are medical pandemics. Bariatric surgery, in the form of Roux-en-Y gastric bypass or sleeve gastrectomy (SG), is currently the most effective and lasting treatment for obesity and related comorbidities^1,2^. For a majority of patients, remission is durable and lasts for years after surgery^1,3^. Two changes consistently observed following bariatric surgery are increased levels of GLP-1, a circulating incretin hormone, and changes in the systemic repertoire of bile acids^4^. Bile acids are cholesterol-derived metabolites that play crucial roles in host metabolism by acting as detergents that aid in the absorption of lipids and vitamins, and as ligands for host receptors^5^. Bile acids have been implicated in post-SG therapeutic benefits due to their ability to mediate signaling through the G protein-coupled bile acid receptor (GPBAR1, also known as TGR5) ^6^ and the farnesoid X receptor (FXR)^7^. Thus far, research efforts have focused on overall changes in the total bile acid pool or in levels of conjugated or unconjugated bile acids in circulating blood^4,8^. Individual bile acids, however, have different binding affinities for nuclear hormone receptors (NhRs) and GPCRs, and thus unique abilities to modulate glucose homeostasis, lipid accumulation, and energy expenditure^5,9^. It is not sufficient, therefore, to limit analyses to whole classes of bile acids. Moreover, local activation of receptors in the intestine can affect global metabolic outcomes. In particular, GLP-1 is secreted post-prandially by enteroendocrine L cells in the lower intestine in response to activation of TGR5, a G-protein coupled receptor with a primary role in energy metabolism^10^. GLP-1 directly stimulates pancreatic insulin release, and both hormones then regulate metabolism systemically ^11^. In this work, we sought to identify specific naturally occurring bile acids whose levels were increased in the gut following SG and then to investigate the role of these compounds as potential TGR5 ligands and thus regulators of glucose metabolism. We identified one bile acid, cholic acid-7-sulfate (CA7S), whose levels were increased in cecal contents and feces of mice and humans, respectively, after SG. We then determined that CA7S is a gut-restricted TGR5 agonist and GLP-1 secretagogue that increases acute glucose tolerance when administered to diet-induced obese mice.

## Results

### Intestinal bile acid profiling reveals increased CA7S in mice and humans post-SG

Owing to robust post-surgical metabolic benefits and favorable side-effect profile, SG is the most common bariatric surgery performed in the US^12^. Rodent SG models mimic the positive metabolic outcomes observed in humans and are thus suitable for studying post-surgical outcomes^13^. In this study, SG or sham surgery was performed on insulin-resistant, diet-induced obese (DIO) mice (Fig. 1a). SG mice displayed improved glucose tolerance and insulin sensitivity 4-5 weeks post-surgery compared to shams (Fig. 1b-e). Mice were euthanized six weeks post-surgery and their tissues were harvested. Consistent with studies involving human patients^4^, we observed an increase in circulating GLP-1 in SG mice (Fig. 1f). Individual bile acids that are known agonists of TGR5 have been shown to induce GLP-1 secretion in lower-intestinal L cells^4,10^. We therefore quantified individual bile acids in cecal contents of SG and sham mice using Ultra-high Performance Liquid Chromatography-Mass Spectrometry (UPLC-MS) (Fig. 2a). We observed a significant increase in only one bile acid in cecal contents of SG mice. Based on its mass, this compound appeared to be a monosulfated metabolite of a trihydroxy bile acid. We purified this compound from pooled extracts of post-SG cecal contents using mass spectrometry-guided semi-preparative HPLC. Using NMR spectroscopy, we then identified this compound as cholic acid-7-sulfate (CA7S) (Fig. 2b,c, Supplementary figs. 1,2). This molecule is a sulfated metabolite of cholic acid (CA), which is an abundant primary bile acid in both mice and humans. Sulfation of bile acids predominantly occurs in the liver^14^. Consistent with this observation, we found increased levels of CA7S in the liver of SG mice (Fig. 2d). While previous studies observed an increase in the total circulating serum bile acid concentration in patients post-bariatric surgery^8^, we did not note a difference in total gut bile acid levels as measured in cecal contents of post-SG compared to post-sham mice (Fig. 2b). Notably, CA7S was the only bile acid detected whose levels were significantly higher in SG mouse cecal contents and livers (Supplementary figs. 3, 4).

**Figure 1.**
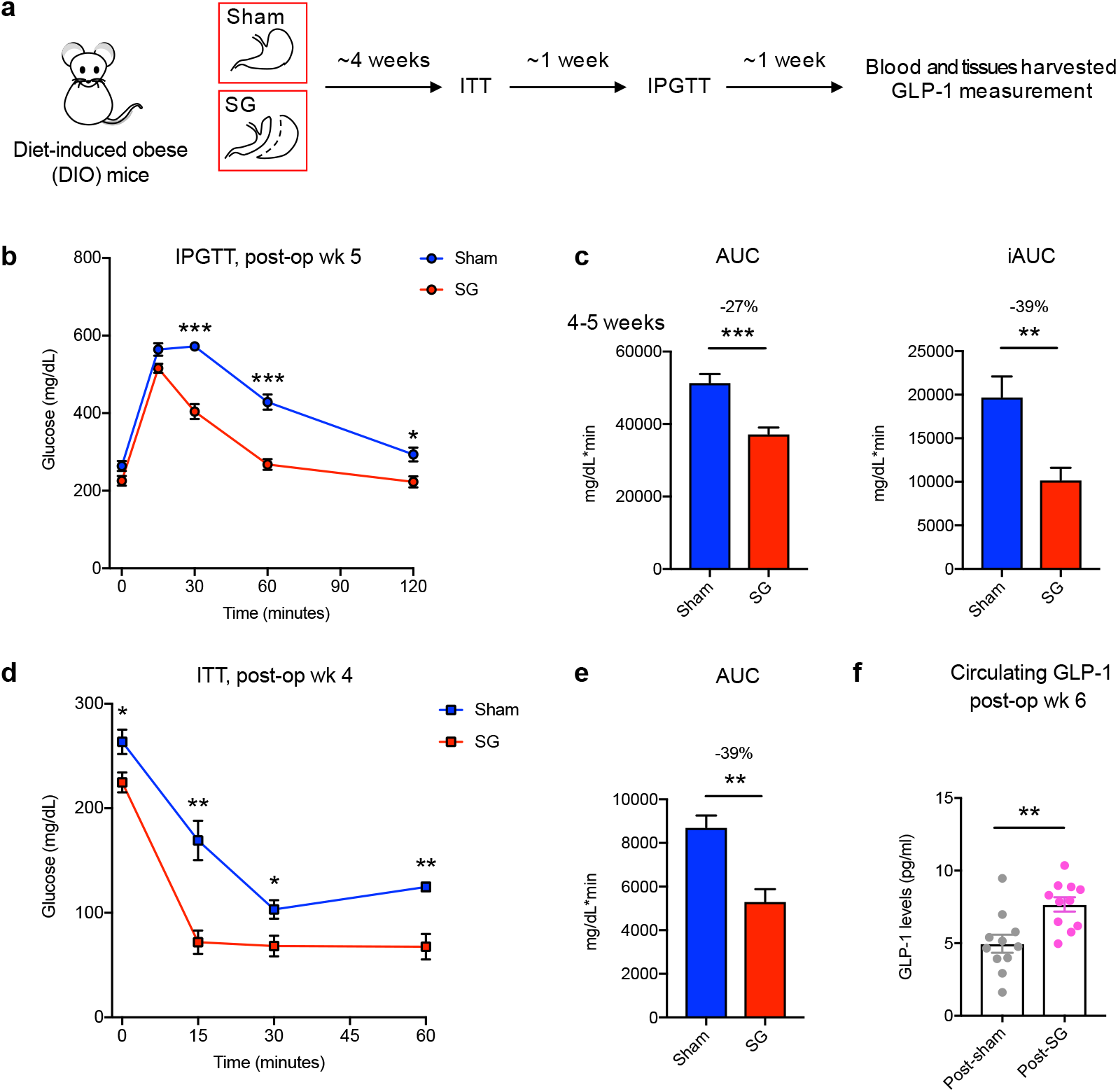
DIO mice display improved glucose tolerance and insulin sensitivity following SG. **a**, Schematic of surgical interventions and post-operative assessments. Sleeve gastrectomy (SG) or sham surgery was performed on diet-induced obese (DIO) mice, followed by an insulin tolerance test (ITT) ~4 weeks post-op and then intraperitoneal glucose tolerance test (IPGTT) ~5 weeks post-op. Blood and tissues were harvested ~6 weeks post-op. **b**, Glycemic curves during IPGTT (SG, n=7; sham, n=6, 30 min \*\*\**p*=5.91×10^−6^, 60 min \*\*\**p*=4.34×10^−5^, 120 min \**p*=0.012, Student’s t test). **c**, Corresponding area under the blood glucose curve (AUC) and incremental area under the curve (iAUC) were reduced in SG-compared to sham-operated mice. (AUC \*\*\**p*= 0.0004, iAUC \*\**p*= 0.006, Student’s t test). **d**, Glycemic curves during ITT (SG, n=4; sham, n=4, 0 min \**p*=0.042, 15 min \*\**p*=0.0043, 30 min \**p*=0.038, 60 min \*\**p*=0.0045, Student’s t test). **e**, Corresponding area under the blood glucose curve (AUC) was reduced in SG-compared to sham-operated mice (AUC \*\**p*= 0.005, Student’s t test). **f**, GLP-1 levels were increased in mice post-SG compared to post-sham (n=11 per group, *\**\**p*=0.003, Welch’s t test). All data are presented as mean ± SEM.

**Figure 2.**
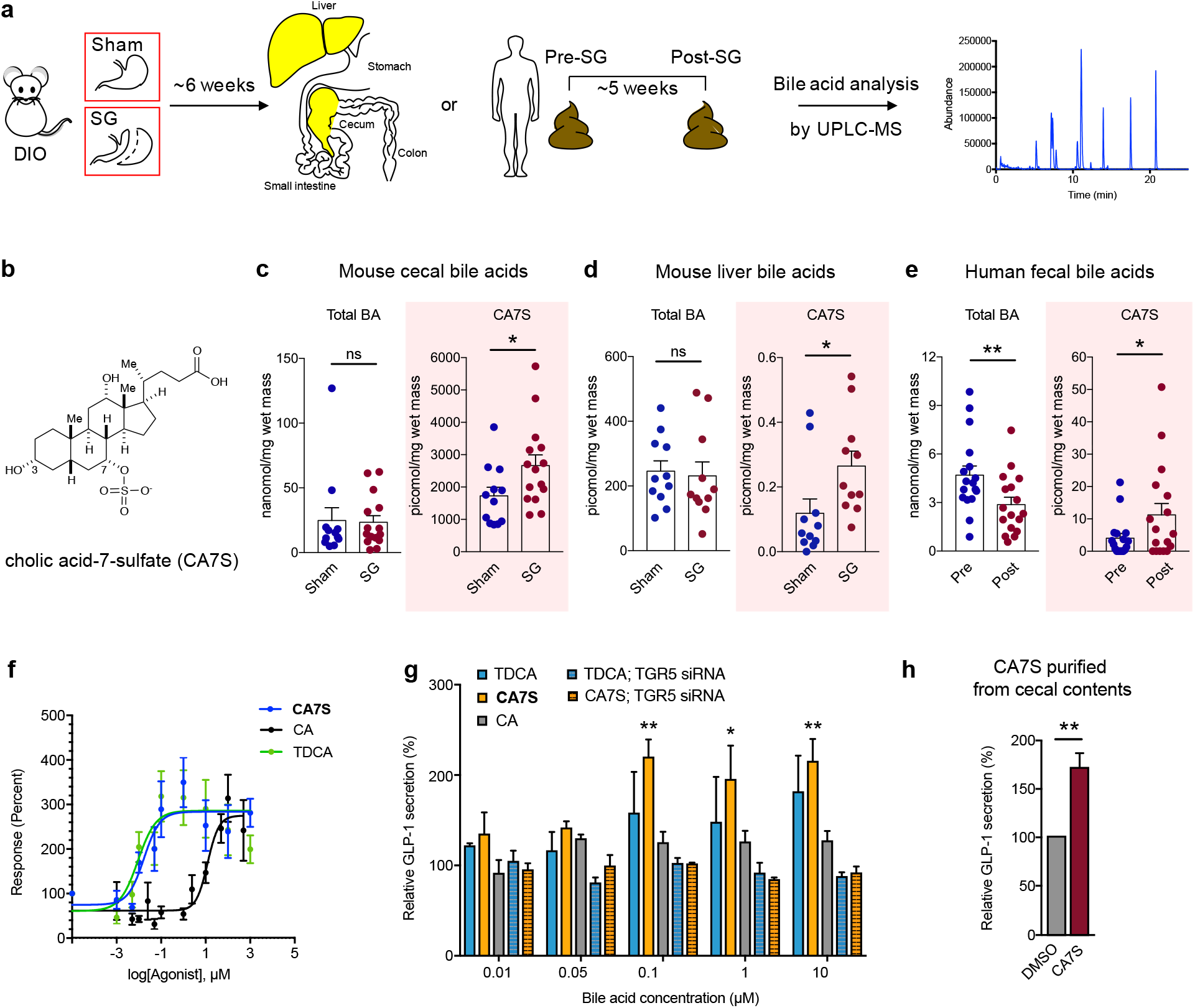
Cholic acid-7-sulfate (CA7S), a bile acid metabolite increased in mice and humans following SG, is a TGR5 agonist and induces GLP-1 secretion in vitro. **a**, Schematic of sample collection followed by bile acid profiling using UPLC-MS. For mice, livers and cecal contents were collected from sham or SG mice 6 weeks post-op. For humans, fecal samples were collected pre-op and ~5 weeks post-op (median 36 days after surgery). **b**, Structure of CA7S. **c**, CA7S was increased in cecal contents of SG mice, while total bile acid concentrations did not differ between SG and sham mice (sham, n=12, SG, n=15, *\*p*=0.037, *p*=0.9, ns=not significant, Welch’s t test). Note that 1 picomol bile acid/mg wet mass is approximately equivalent to 1 μM. **d**, CA7S was increased in livers of SG mice (n=11 per group, *\*p*=0.033, *p*=0.546, ns=not significant, Welch’s t test). **e**, CA7S in human feces was increased post-SG compared to pre-surgery (n=17 patients, *\*p*=0.015, *\*\*p*=0.001, paired t test). **f**, Dose response curves for human TGR5 activation in HEK293T cells overexpressing human TGR5 for CA7S, TDCA, CA (≥3 biological replicates per condition). **g**, CA7S-induced secretion of GLP-1 in NCI-H716 cells compared to both CA and the known TGR5 agonist, TDCA. siRNA-mediated knockdown of TGR5 abolished GLP-1 secretion (≥3 biological replicates per condition, 0.1μM \*\**p*=0.008, 1μM \**p*=0.030, 10μM \*\**p*=0.002, one-way ANOVA followed by Dunnett’s multiple comparisons test). **h**, CA7S (500 μM) purified from SG mouse cecal contents induced secretion of GLP-1 in NCI-H716 cells compared to DMSO control (*\**\**p*=0.001, Welch’s t test). All data are presented as mean ± SEM.

To determine whether CA7S concentrations were also higher in humans after surgery, we quantified bile acids in stool from patients who had undergone SG. Remarkably, even though total fecal bile acid levels were decreased in patients post-SG compared to their pre-surgery levels, fecal CA7S levels were significantly increased (Fig. 2e, Supplementary fig. 5). To our knowledge, this is the first report of a specific bile acid metabolite that is significantly increased following SG in both mice and human subjects. Similar to DIO mouse cecal contents, there was no significant difference in total bile acid concentration in feces before or after SG in these patients (Fig. 2e, Supplementary fig. 5).

### CA7S activates TGR5 and induces GLP-1 secretion in vitro

We next sought to determine whether CA7S can activate TGR5 and thereby induce GLP-1 secretion from L cells. Previous work has shown that sulfation of both natural bile acids and synthetic analogs significantly alters the TGR5 agonistic activity of these compounds^15^. We therefore hypothesized that CA7S might possess altered TGR5 agonism compared to CA. While sulfation at C3 abolishes the TGR5 agonist activity of lithocholic acid (EC_50_ values of 0.58 μM and >100 μM, respectively), replacement of the C24-carboxylic acid with a C24-sulfate lowers the EC_50_ of CA, chenodeoxycholic acid, and ursodeoxycholic acid by an order of magnitude in each case^15^. Based on these results, we hypothesized that the C7-sulfated version of CA, CA7S, might possess significantly different TGR5 agonistic activities than CA, which is a weak agonist of TGR5 (reported EC_50_ value of 13.6 uM)^15^. We examined the activation of human TGR5 in human embryonic kidney cells (HEK293T) by CA7S, CA, or tauro-deoxycholic acid (TDCA), which is a naturally occurring bile acid and potent TGR5 agonist^16^. We found that CA7S activated human TGR5 in a dose-dependent manner and to a similar extent as TDCA. CA7S also displayed a lower EC_50_ (0.17 μM) than CA (12.22 μM) (Fig. 2f).

TDCA is currently one of the most potent, naturally occurring GLP-1 secretagogues known^16^. We observed that CA7S induced GLP-1 secretion in human intestinal L cells (NCI-H716) to a similar degree as TDCA in a dose-dependent manner, while CA had no effect on GLP-1 secretion (Fig. 2g and Supplementary fig. 6a). CA7S extracted and purified directly from cecal contents of SG mice also induced GLP-1 secretion in vitro (Fig. 2h). Furthermore, siRNA-mediated knockdown of TGR5 abolished both CA7S- and TDCA-mediated secretion of GLP-1 (Fig. 2g, Supplementary fig. 6a,b). This result indicates that induction of GLP-1 secretion by CA7S requires TGR5. TGR5 agonism also results in elevated intracellular calcium levels^17^. Consistent with this previous finding, we observed a dose-dependent increase in calcium levels in NCI-H716 cells treated with CA7S (Supplementary fig. 6c). Taken together, our results demonstrate that CA7S, a naturally occurring bile acid metabolite, is a potent TGR5 agonist and GLP-1 secretagogue.

### Acute enteral CA7S administration induces GLP-1 expression and reduces blood glucose in vivo

We next evaluated the acute anti-diabetic effects of CA7S in vivo. DIO mice were treated with either CA7S or PBS via duodenal and rectal catheters (Fig. 3a). Administration of 1 mg of CA7S resulted in an average of 2500 pmol/mg wet mass of CA7S in cecal contents, a concentration similar to observed post-SG levels (Fig. 2b, 3b, Table 1). Consistent with our in vitro studies, CA7S-treated mice displayed increased systemic GLP-1 levels compared to PBS-treated mice within 15 minutes (Fig. 3c). Moreover, CA7S-treated mice exhibited reduced blood glucose levels and increased insulin levels compared to PBS-treated mice (Fig. 3d,e, Supplementary fig. 6d). GLP-1-producing enteroendocrine L cells are enriched in the distal compared to the proximal gut^18,19^. Consistent with this finding, we observed that TGR5 expression was increased in the colon, but not the terminal ileum, of CA7S-treated mice (Fig. 3f). These results suggest that in an acute setting, distal action of CA7S in the GI tract induces systemic glucose clearance and thus ameliorates hyperglycemia.

**Table 1.** Cholic acid-7-sulfate concentration in indicated tissues and blood.

| Treatment | Tissue/blood | CA7S concentration (mean ± SEM) |
| --- | --- | --- |
| DIO mice; sham surgery | Cecum | 1726 ± 267 pmol/mg |
| | Liver | 0.12 ± 0.04 pmol/mg |
|  | Portal vein | n.d. |
|  | Systemic blood | n.d. |
| DIO mice; sleeve<br>gastrectomy | Cecum | 2661 ± 331 pmol/mg |
|  | Liver | 0.27 ± 0.04 pmol/mg |
|  | Portal vein | n.d. |
|  | Systemic blood | n.d. |
| DIO mice; enteral PBS | Cecum | 161.1 ± 46.4 pmol/mg |
|  | Portal vein | 0.07 ± 0.06 pmol/mg |
|  | Systemic blood | n.d. |
| DIO mice; enteral CA7S | Cecum | 2577 ± 185 pmol/mg |
|  | Portal vein | 6.13 ± 2.11 pmol/mg |
|  | Systemic blood | 0.5 ± 0.2 pmol/μl |
| DIO mice; PBS gavage | Cecum | 947 ± 349 pmol/mg |
|  | Portal vein | n.d. |
|  | Systemic blood | n.d. |
| DIO mice; CA7S gavage | Cecum | 14345 ± 1451 pmol/μl |
|  | Portal vein | 13.2 ± 7.7 pmol/μl |
|  | Systemic blood | n.d. |

n.d. not detected, all data are presented as mean ± SEM.

**Figure 3.**
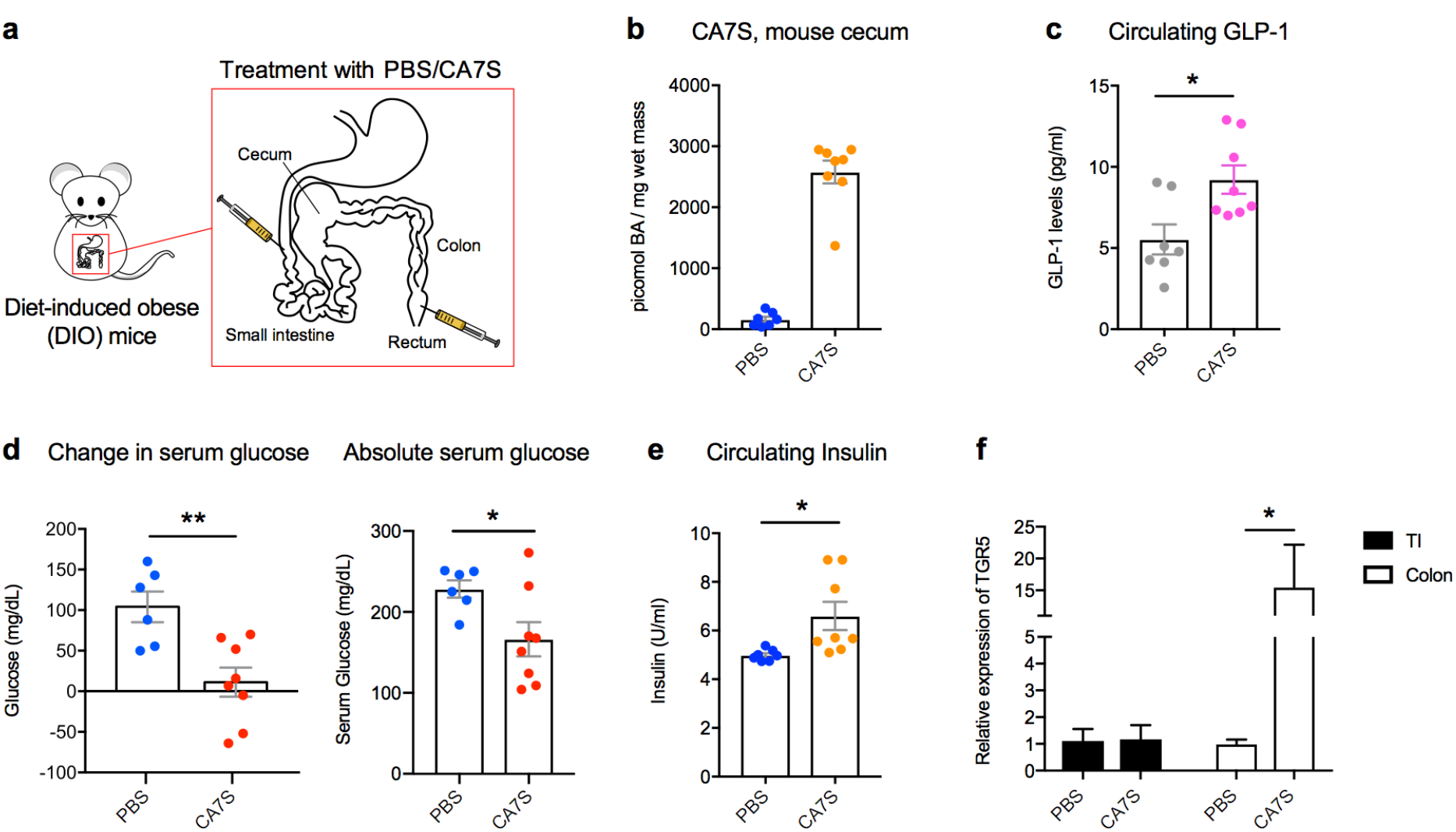
Acute CA7S administration induces GLP-1 and reduces serum glucose levels in vivo. **a**, Schematic of acute treatment wherein anesthetized DIO mice were treated with PBS or CA7S via duodenal and rectal catheters. **b**, Concentration of CA7S in mouse cecum 15 minutes after treatment with PBS or CA7S (PBS, n=7; CA7S, n=8 mice per group). **c**-**e**, CA7S-treated mice displayed increased GLP-1 (**c**), reduced blood glucose levels (**d**), and increased blood insulin levels (**e**) compared to PBS-treated mice. (For **c**and **e**, PBS, n=7; CA7S, n=8 mice per group, (**c**) \**p*=0.012, (**e**) \**p*=0.023, Welch’s t test. For **d**, PBS, n=6; CA7S, n=8 mice per group, \*\**p*=0.0043, \**p*=0.033, Student’s t test). **f**, CA7S treatment in the intestine induced TGR5 expression in the colon but not in the terminal ileum (TI) (\**p*=0.035, Welch’s t test). All data are presented as mean ± SEM.

### Oral CA7S administration increases glucose tolerance in vivo

To further study the anti-diabetic effects of CA7S, DIO mice were orally gavaged with CA7S at a dose of 100 mg/kg (Fig. 4a). Analysis of cecal contents 5 hours post-gavage showed an accumulation of 15,000 picomol/mg wet mass of CA7S (mean value, Fig. 4b), a concentration that is within an order of magnitude of the mean amount measured in post-SG mice. These data indicate that we had administered a physiologically relevant concentration of this metabolite. Systemic levels of GLP-1 were increased in CA7S-gavaged mice compared to PBS-treated mice 5 hours post-treatment (Fig. 4c). This result is consistent with the findings from enteral administration and demonstrates that oral CA7S treatment can increase circulating GLP-1 for several hours.

**Figure 4.**
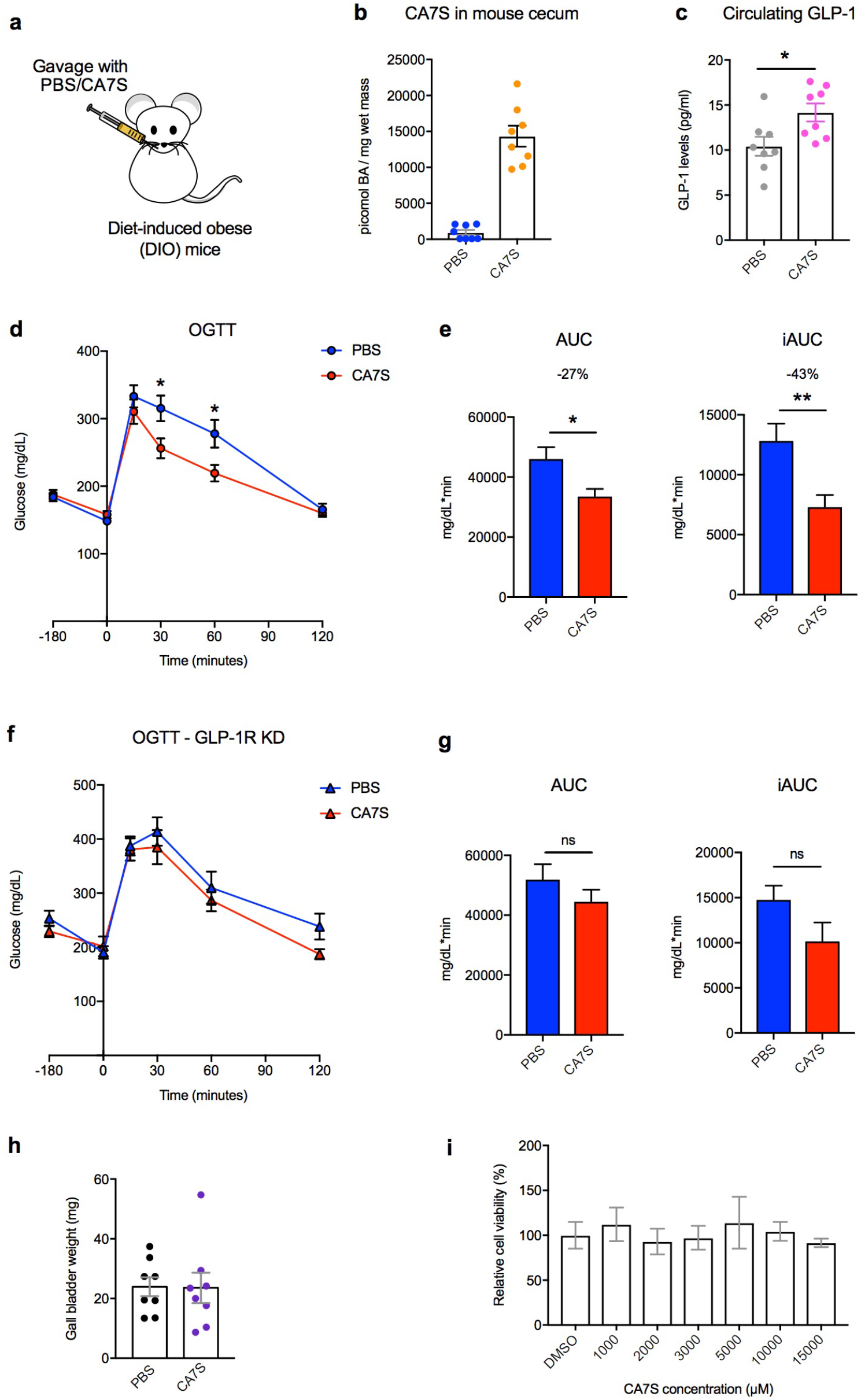
CA7S gavage induces GLP-1 and improves glucose tolerance in vivo via GLP-1R. **a**, Schematic of treatment of DIO mice with either PBS or CA7S via oral gavage. **b**, Concentration of CA7S in mouse cecum 5 hours after PBS or CA7S gavage. **c**, CA7S mice displayed increased GLP-1 levels 5 hours post-gavage (For **b** and **c**, n=8 mice per group, \**p*=0.021, Welch’s t test). **d**,**e**, DIO mice treated with CA7S (100 mg/kg) displayed increased glucose tolerance compared to vehicle-treated mice 3 hours post-gavage as determined by an oral glucose tolerance test (OGTT) (n=11 mice per group). **d**, Glycemic curves during OGTT (\**p*=0.023, Student’s t test). **e**, Corresponding blood glucose AUC and iAUC were significantly reduced in CA7S-treated mice (AUC \**p*=0.011, iAUC \*\**p*=0.004, Student’s t test). **f**,**g**, On day 3 after treatment with lentiviral shRNA targeting GLP-1 receptor (GLP-1R), CA7S (100 mg/kg) or PBS was administered, and 3 hours later, an OGTT was performed (n=7 mice per group). **f**, Glycemic curves during OGTT (ns = not significant, 30 min *p*=0.49, 60 min *p*=0.53, 120 min *p*=0.07, Student’s t test). **g**, Corresponding blood glucose AUC and iAUC were not significantly different in CA7S- or PBS-treated mice in which the GLP-1R had been knocked down (ns = not significant, AUC *p*=0.266, iAUC *p*=0.1, Student’s t test). **h**, CA7S treatment does not induce gallbladder filling in mice (n=8 mice per group, not significant, *p*=0.95, Welch’s t test). **i**, Percentage cell viability upon treatment of Caco-2 cells with CA7S in vitro (≥3 biological replicates per condition, not significant, *p*≥0.97 one-way ANOVA followed by Dunnett’s multiple comparisons test). All data are presented as mean ± SEM.

We then determined the effect of CA7S on glucose tolerance over time using an oral glucose tolerance test (OGTT). DIO mice were gavaged with CA7S (100 mg/kg) or PBS and then administered an oral glucose bolus 3 hours later. CA7S treatment resulted in an increased rate of blood glucose clearance (Fig. 4d). Moreover, the total and incremental areas under the glucose versus time curves (AUC and iAUC) were significantly decreased in CA7S- compared to vehicle-treated mice (Fig. 4e). These results demonstrate that CA7S increases blood glucose clearance following oral glucose challenge, a clinically relevant test used in the diagnosis of diabetes.

### Anti-diabetic effects of CA7S are largely dependent on GLP-1

Because CA7S is a potent inducer of GLP-1 secretion, we sought to determine if the acute anti-diabetic effects of CA7S are dependent on GLP-1. To investigate this question, we performed lentiviral shRNA-mediated knockdown of the GLP-1 receptor (GLP-1R) in vivo. DIO mice were injected intraperitoneally with 5×10^5^ shRNA lentiviral particles targeting GLP-1R. At day 3 post-injection, expression of GLP-1R in the small intestine, heart, and stomach was significantly reduced, and importantly, the expression of GLP-1R was undetectable in the pancreas (Supplementary fig. 7a). Mice were then gavaged with CA7S (100 mg/kg) or PBS and subjected to an OGTT 3 hours post-gavage. We observed that while the glycemic curve and AUCs for the CA7S-treated animals were lower than those of the PBS-treated mice, there were no longer significant differences in these metrics in the absence of GLP-1R. These data suggest that in an acute setting, the blood glucose clearing-effects of CA7S are largely dependent on the action of GLP-1 (Fig. 4f,g).

### CA7S is gut-restricted, non-toxic, and does not induce gallbladder filling

While high concentrations of CA7S were observed in the intestine, this metabolite was undetectable in both circulating and portal venous blood from SG and sham-operated mice (Table 1). This result suggests that CA7S is neither recycled via enterohepatic circulation nor absorbed into systemic circulation. In contrast, the known endogenous TGR5 agonists TDCA and DCA are found in systemic circulation in mice^9^. Introduction of CA7S via enteral administration or oral gavage resulted in only minor amounts in circulating and portal venous blood (Table 1). Our findings are consistent with previous observations that sulfated bile acids, in particular 7α-sulfated bile acids, are poorly absorbed in the intestine^14^.

Notably, while synthetic TGR5 agonists ameliorate diabetic phenotypes^20^, their use as therapeutics is hampered by significant side effects resulting from their absorption into circulation ^20,21^. In particular, systemically absorbed TGR5 agonists reduce bile secretion and induce gallbladder filling, conditions that cause bile stasis^10,21^. We did not observe any change in gallbladder weight in DIO mice 5 hours after oral gavage of CA7S compared to PBS control treatment (Fig. 4h). This result suggests that, in an acute setting, CA7S does not display one of the major side-effects of non-gut-restricted TGR5 agonists. Finally, CA7S does not affect the viability of Caco-2 cells at concentrations up to 15 mM (Fig. 4i), indicating that this metabolite is not toxic to human intestinal cells at physiologically relevant concentrations.

## Discussion

By quantifying changes in individual bile acids, we found that the sulfated metabolite CA7S is increased in both the mouse GI tract and human feces following SG. We then found that CA7S is a potent TGR5 agonist that induces the secretion of the incretin hormone, GLP-1 from enteroendocrine L cells both in vitro and in vivo. In acute settings, we observed that CA7S exhibits anti-diabetic effects, including decreasing blood glucose levels and increasing glucose tolerance in insulin-resistant mice. Unlike known endogenous TGR5 agonists, CA7S is not absorbed into portal or systemic circulation. CA7S had previously been regarded as a metabolic waste product produced by the liver – a sulfated form of cholic acid that was tagged for excretion in feces^14^. To our knowledge, no role for CA7S as a ligand for a GPCR or NhR has been reported. Taken together, our data indicate that CA7S is a signaling molecule that binds to TGR5 exclusively in the gut and thereby increases systemic levels of GLP-1.

Owing to the significant off-target effects caused by TGR5 agonists absorbed into circulation, it has been suggested that an improved TGR5-based therapeutic for T2D would specifically activate intestinal TGR5^21^. This GPCR is expressed both apically and basolaterally on L cells^16^. TDCA has been shown to activate TGR5 and induce GLP-1 secretion when applied to L cells from both apical and basolateral directions ^16^. These findings suggest that development of a gut-restricted TGR5 agonist could be effective in the treatment of T2D. In the setting of acute administration, we found that CA7S does not induce gallbladder filling as has been reported for synthetic TGR5 agonists that enter circulating blood. Furthermore, CA7S is not toxic to human intestinal cells and is stable at physiological pHs (Supplementary fig. 7b). Additional studies are required to assess the long-term effects of CA7S on glucose tolerance and insulin sensitivity in vivo. Nonetheless, as a result of its beneficial acute metabolic effects, gut restriction, and low toxicity, CA7S could be a candidate for the development of a T2D therapeutic.

The role that CA7S plays in the metabolic changes observed post-SG remains unknown. Due to the redundancy of sulfotransferases (SULTs), the enzymes responsible for CA7S synthesis in the the mouse liver^22^, knock-down of endogenous CA7S levels may be a challenging endeavor. If this obstacle could be overcome, performing SG on CA7S-depleted mice would help to reveal the contribution of this molecule to the metabolic effects of the surgery.

More broadly, prior to this work, there were no known individual metabolites whose levels were altered by bariatric surgery that could increase blood glucose clearance. Through the identification and study of CA7S, we have uncovered a metabolite that, while restricted to the GI tract, can improve global glucose regulation.

## Online Methods

### Animals

Diet induced obese, male, C57Bl/6J mice were purchased from Jackson Laboratory (Bar Harbor, ME) at 11-16 weeks of age. They were housed under standard conditions in a climate-controlled environment with 12 hour light and dark cycles and reared on a high fat diet (HFD, 60% Kcal fat; RD12492; Research Diets, NJ). They were allowed to acclimate for at least 1 week prior to undergoing any procedures. All animals were cared for according to guidelines set forth by the American Association for Laboratory Animal Science. All procedures were approved by the Institutional Animal Care and Use Committee.

### Sleeve gastrectomy (SG) and sham procedures

11-week-old DIO mice were purchased and housed as described above. Mice were weight-matched into two groups and either underwent SG or Sham operation. SG was performed through a 1.5 cm midline laparotomy under isoflurane anesthesia. The stomach was gently dissected free from its surrounding attachments, the vessels between the spleen and stomach (short gastric vessels) were divided, and a tubular stomach was created by removing 80% of the glandular and 100% of the non-glandular stomach with a linear-cutting surgical stapler. Sham operation consisted of a similar laparotomy, stomach dissection, ligation of short gastric vessels, and manipulation of the stomach along the staple line equivalent. Mice were then individually housed thereafter to allow for monitoring of food intake, weight, and behavior. SG and Sham mice were maintained on Recovery Gel Diet (Clear H_2_O, Westbrook, ME) from 1 day prior through 6 days after surgery and then were restarted on HFD on the morning of post-operative day (POD) 7. Mice were sacrificed 5-7 weeks post-surgery.

### Functional glucose testing

After a 4 hour fast (8 am to noon), intraperitoneal glucose tolerance testing (IPGTT) and insulin tolerance testing (ITT) were performed at post-operative week 4 and 5, respectively. During IPGTT, mice received 2 mg/g of intraperitoneal D-Glucose (Sigma-Aldrich, St. Louis, MO) and serum glucose levels were measured from the tail vein at 15, 30, 60, and 120 min with a OneTouch Glucometer (Life technologies, San Diego, CA). ITT was performed by intraperitoneal instillation of 0.6u/kg of regular human insulin (Eli Lily and Company, Indianapolis, IN) and measurement of serum glucose levels at 15, 30, and 60 min. Baseline glucose was measured for each set prior to medication administration.

### Body weight and food intake measurements

Mice were individually housed and weighed daily for the first post-operative week and then twice weekly until sacrifice. Food intake was measured twice weekly and daily food intake was calculated by averaging the grams eaten per day over the preceding days. Note that food intake measurements were started on POD 10 as animals were transitioned from Gel Diet to high fat diet on the morning of POD 7.

### Bile acid analysis

Bile acid analyses were performed using a previously reported method^23^.

#### Reagents

Stock solutions of all bile acids were prepared by dissolving the compounds in molecular biology grade DMSO (VWR International, Radnor, PA). These solutions were used to establish standard curves. CA7S was purchased from (Caymen Chemicals, Ann Arbor, MI. Cat. No. 9002532). Glyocholic acid (GCA) (Sigma) was used as the internal standard for measurements in mouse tissues. HPLC grade solvents were used for preparing and running UPLC-MS samples.

#### Extraction

Cecal, liver, and human fecal samples (approximately 50 mg each) and mouse portal veins were pre-weighed in lysis tubes containing ceramic beads (Precellys lysing kit tough micro-organism lysing VK05 tubes for cecal, fecal samples, and portal veins; tissue homogenizing CKMix tubes for liver samples; Bertin technologies, Montigny-le-Bretonneux, France). 400μL of methanol containing 10 μM internal standard (GCA) was added and the tubes were homogenized in a MagNA Lyser (6000 speed for 90 s*2, 7000 speed for 60 s). 50 μl of mouse serum was collected in Eppendorf tubes, followed by addition of 50 μL of methanol containing 10 μM internal standard (GCA). After vortexing for 1 min, the sample was cooled to −20 °C for 20 min. All methanol-extracted tissue samples were centrifuged at 4 °C for 30 min at 15,000 rpm. The supernatant was diluted 1:1 in 50% methanol/water and centrifuged again at 4 °C for 30 min at 15000 rpm. The supernatant was transferred into mass spec vials and injected into the UPLC-MS.

#### UPLC-MS analysis

Samples were injected onto a Phenomenex 1.7 μm, C18 100 Å, 100 × 21 mm LC column at room temperature and eluted using a 30 min gradient of 75% A to 100% B (A = water + 0.05% formic acid; B = acetone + 0.05% formic acid) at a flow rate of 0.350 mL/min. Samples were analyzed using an Agilent Technologies 1290 Infinity II UPLC system coupled online to an Agilent Technologies 6120 Quadrupole LC/MS spectrometer in negative electrospray mode with a scan range of 350–550 m/z (MSD settings: fragmentor - 250, gain - 3.00, threshold - 150, Step size - 0.10, speed (u/sec) - 743). Capillary voltage was 4500 V, drying gas temperature was 300°C, and drying gas flow was 3 L/min. Analytes were identified according to their mass and retention time. For quantification of the analytes, standard curves were obtained using known bile acids, and then each analyte was quantified based on the standard curve and normalized based on the internal standard. The limits of detection for individual bile acids were determined using commercially available standards solubilized in 1:1 MeOH/water and are as follows: CA7S, 0.05 picomol/μL; α/βMCA, 0.03 picomol/μL; Tα/βMCA, 0.01 picomol/μL; CA, 0.04 picomol/μL; TCA, 0.01 picomol/μL; UDCA, 0.04 picomol/μL; TUDCA, 0.01; CDCA, 0.04 picmol/μL; TCDCA, 0.01 picomol/μL; LCA, 0.03 picomol/μL; isoLCA, 0.07 picomol/μL; 3-oxo-LCA, 0.05 picomol/μL; DCA, 0.04 picomol/μL; 3-oxo-CA, 0.04 picomol/μL; 3-oxo-CDCA, 0.4 picomol/μL; 7-oxo-CDCA, 0.03 picomol/μL. Note that CA7S and cholic acid-3-sulfate can be distinguished based on retention time using this UPLC-MS method.

#### Purification of CA7S

Extracted cecal contents from 11 SG mice (same shown in Fig. 1) were pooled to provide sufficient material for purification. Pooled extract was purified via MS-guided HPLC of *m/z* 487 using a Luna RP C18 semi-preparative column and water and acetonitrile with 0.1% formic acid as an additive.

#### NMR spectroscopy

CA7S and purified *m/z* 487 (<1 mg) were dissolved in 250 μL DMSO-d6. Nuclear magnetic resonance (NMR) spectra were acquired on a Varian INOVA 500 MHz and are referenced internally according to residual solvent signals (DMSO to 2.50, HOD to 3.33).

### Cell culture

NCI-H716 cells and Caco-2 cells were obtained from American Type Culture Collection (Manassas, VA). HEK-293T cells were a kind gift from the Blacklow lab (BCMP, HMS). Caco-2 and HEK-293T cells were maintained in Minimum Essential Medium (MEM) with GlutaMAX and Earle’s Salts (Gibco, Life Technologies, UK). NCI-H716 cells were maintained in RPMI 1640 with L-glutamine (GenClone, San Diego, CA). All cell culture media were supplemented with 10% fetal bovine serum (FBS), 100 units/ml penicillin, and 100 μg/ml streptomycin (GenClone). Cells were grown in FBS- and antibiotic-supplemented ‘complete’ media at 37 °C in an atmosphere of 5% CO_2_.

### In vitro bile acid treatments

NCI-H716 cells were seeded in cell culture plates coated with Matrigel (Corning, NY. Cat. No. 356234) diluted in Hank’s Balanced Salt Solution (HBSS, Gibco) according to manufacturer’s instructions. The cells were allowed to grow for 2 days in complete RPMI media. On the day of the treatment, cells were rinsed gently with low serum (0.5% FBS) RPMI 1640 medium without antibiotics. Bile acids cholic acid-7-sulfate (CA7S), cholic acid (CA) (Sigma) and taurodeoxycholic acid (TDCA) (Sigma) were diluted in dimethyl sulfoxide (DMSO, VWR International) and added to cells in the low serum media (0.5% FBS, RPMI 1640) without antibiotics. The concentration of DMSO was kept constant throughout the treatments and used as a negative control. Cells were incubated at 37°C in an atmosphere of 5% CO_2_ for 2 hours. After the incubation period, cell culture media was collected in Eppendorf tubes containing 1% trifluoroacetic acid (TFA, Sigma) in sterile purified water (GenClone) to make a final TFA concentration of 0.1% and frozen at −80 °C for further GLP-1 measurements. Cells on cell culture plates were placed on ice and gently washed with PBS (GenClone). Cells used for GLP-1 measurements were treated with ice-cold cell lysis solution of 1% TFA, 1N hydrochloric acid, 5% formic acid, and 1% NaCl (all from Sigma), scraped off of the Matrigel coating, and collected in lysing tubes with ceramic beads (Precellys lysing kit tough micro-organism lysing VK05 tubes). For calcium measurements, PBS was added to cells, and were collected in lysing tubes containing ceramic beads (Precellys lysing kit tough micro-organism lysing VK05 tubes). Cells were thereafter lysed in a MagNA Lyser and stored at −80 °C for further analysis. Cells used for RNA extraction were treated with TRIzol (Ambion, Life Technologies, Thermo Fisher Scientific, Waltham, MA) and stored at −80 °C for further analysis.

### GLP-1 and Insulin measurements

Total GLP-1 peptide measurements were performed using the GLP-1 EIA Kit (Sigma, Cat. No. RAB0201) and total insulin levels were measured using the Mouse Ins1/Insulin-1 ELISA kit (Sigma, Cat. No. RAB0817) according to manufacturer’s instructions. Mouse serum samples, NCI-H716 cell lysates, and cell culture media samples were stored at −80 °C and thawed on ice prior to performance ELISA assay. 20 μl of mouse serum samples were used directly in the GLP-1 ELISA assay, while 50 μl of mouse serum samples were used directly in the Insulin ELISA assay. Cell culture media were centrifuged at 12000 rpm, and the supernatant was directly used in the GLP-1 ELISA assay. Cell lysates were subjected to peptide purification using Sep Pak C18 Classic columns (Waters Corporation, Milford, MA). The column was pretreated with a solution of 0.1% TFA in 80% isopropyl alcohol (EMD Millipore) and equilibrated with 0.1% TFA in water. Cell lysates were loaded onto the column and washed with 0.1% TFA in 80% isopropyl alcohol. The peptides were eluted in 0.1% TFA in water. The eluate was concentrated by drying under vacuum and resuspended in 0.1% TFA in water. Water was used as ‘blank’ reading for serum GLP-1 ELISA, while 0.1% TFA in water was used as ‘blank’ for cell culture media and purified cell lysate ELISAs. Excess samples were stored at −80°C for later analyses. Total GLP-1 amounts in the cell culture media (secreted) and cell lysates were calculated using a standard curve provided in the EIA kit. Percentage GLP-1 secretion was calculated as follows:

% GLP-1 secretion = total GLP-1 secreted (media) / (total GLP-1 secreted (media) + total GLP-1 in cell lysates) * 100. Relative GLP-1 secretion was calculated compared to DMSO control.

### Plasmids and transient transfections

Human TGR5 was cloned using cDNA from human Caco-2 cells as template and a forward primer with an *EcoRI* restriction-site (5’-CGGAATTCGCACTTGGTCCTTGTGCTCT-3’) and a reverse primer with a *XhoI*-site (5’-GTCTCGAGTTAGTTCAAGTCCAGGTCGA-3’). The PCR product was cloned into the pCDNA 3.1+ plasmid (Promega Corporation, Madison, WI) and transfected at a concentration of 0.4 μg/ml of media. For luciferase reporter assays for TGR5 activation, the pGL4.29[luc2P/CRE/Hygro] plasmid (Promega Corporation), and the pGL4.74[hRluc/CMV] plasmid (Promega Corporation) were used at concentration of 2 μg/ml and 0.05 μg/ml of media respectively. All plasmids were transfected using Opti-MEM (Gibco) and Lipofectamine 2000 (Invitrogen, Life Technologies, Grand Island, NY, USA) according to manufacturer’s instructions. Plasmid transfection were performed in antibiotic-free media (MEM for HEK293T and RPMI for Matrigel-attached NCI-H716 cells) with 10% FBS. After overnight incubation, bile acids were added in complete media and incubated overnight. Cells were harvested the next day for luciferase assay. TGR5 siRNA (Santa Cruz Biotechnology, Dallas, TX) and negative siRNA (Ambion) transfection was performed using Opti-MEM and Lipofectamine 2000 according to manufacturer’s instructions. After siRNA transfection, cells were incubated in antibiotic- and serum-free media (RPMI for Matrigel-attached NCI-H716 cells) for 24 hours. The next day, the media was replaced by complete media and incubated overnight. Bile acids were added 48 hours post-siRNA transfection in complete media and incubated overnight. Cells were harvested the next day for luciferase assay or RNA extraction.

### Luciferase reporter assay

Luminescence was measured using the Dual-Luciferase Reporter Assay System (Promega Corporation) according to manufacturer’s instructions. Cells were washed gently with PBS and lysed in PLB from the kit. Matrigel-attached cells were scraped in PLB. Luminescence was measured using the SpectraMax M5 plate reader (Molecular Devices, San Jose, CA) at the ICCB-Longwood Screening Facility at Harvard Medical School. Luminescence was normalized to *Renilla* luciferase activity and percentage relative luminescence was calculated compared to DMSO control.

### Calcium measurement

CA7S-treated NCI-H716 cells collected in PBS were used to measure intracellular calcium using the Calcium Assay Kit (Fluorometric) (Abcam, UK). Cell lysates were centrifuged at 12000 rpm, and the supernatant was directly used in the calcium assay according to manufacturer’s instructions. Fluorescence was measured using the SpectraMax M5 plate reader (Molecular Devices, San Jose, CA) at the ICCB-Longwood Screening Facility at HMS. Percentage relative fluorescence was calculated compared to DMSO control.

### Cell viability assay

Caco-2 cells were treated with CA7S diluted in DMSO in complete MEM media. The concentration of DMSO was kept constant and used as a negative control. Cells were incubated with CA7S overnight at 37 °C in an atmosphere of 5% CO_2_. The next day, cells were treated with 0.25% trypsin in HBSS (GenClone) for 10 min at 37°C. Cell viability was measured in Countess II automated cell counter (Invitrogen). Percentage relative viability was calculated compared to DMSO control.

### pH stability test

Stability of CA7S in physiological pH’s was determined using the Waters pH stability test. Briefly, buffers of pH 1 (0.1M HCl), pH7.4 (PBS) and pH 9 (a 10 mM solution of ammonium formate adjusted to pH 9 with ammonium hydroxide) (all from Sigma) were prepared. CA7S was incubated in the pH buffers overnight at 37 °C with gentle shaking (50 rpm). The next day, the CA7S solution was diluted in methanol, transferred into mass spec vials and injected into the UPLC-MS.

### RNA extraction and qPCR

Cells frozen in TRIzol (Ambion) were collected in RNase-free Eppendorf tubes and vortexed for 30 seconds. Tissues were collected in Precellys tubes with ceramic beads and TRIzol, followed by homogenization in a MagNA Lyser (Roche, Switzerland). Tubes were kept on ice whenever possible. Chloroform was added (200μl chloroform/1ml TRIzol) and vortexed for 30 seconds. Tubes were centrifuged at 12,000 rpm for 15 min at 4 °C. The clear top layer was transferred to new RNase-free Eppendorf tubes containing 2-propanol and inverted to mix (500μl 2-propanol/1ml TRIzol). Tubes were centrifuged at 12,000 rpm for 10 min at 4 °C. The pellet was washed with 70% EtOH and centrifuged at 14,000rpm for 5 minutes at 4 °C. The RNA pellet was air-dried and resuspended in RNase-free H_2_O (GenClone). cDNA synthesis was performed using the High Capacity cDNA Reverse Transcription Kit (Applied Biosystems, Invitrogen, Foster City, CA). qPCR was performed using the Lightcycler 480 SYBR Green I Mater (Roche, Switzerland) in a 384-well format using a LightCycler 480 System (Roche) at the ICCB-Longwood Screening Facility at Harvard Medical School. Cq values above 45 were considered as not detected (n.d.). The 2^−ΔΔCt^ method was used to calculate the relative change in gene expression. Human *TGR5* gene expression were normalized to the human *HPRT1* (*HGPRT*). Mouse *GLP-1R* gene expression was normalized to 18S. Primer sequences were: human *TGR5*: Forward: 5’-CCTAGGAAGTGCCAGTGCAG-3’, Reverse: 5’-CTTGGGTGGTAGGCAATGCT-3’; human *HGPRT*: Forward: 5’-CCTGGCGTCGTGATTAGTGA-3’, Reverse: 5’ CGAGCAAGACGTTCAGTCCT-3’; mouse *GLP-1R*: Forward: 5’-AGGGCTTGATGGTGGCTATC-3’, Reverse: 5’-GGACACTTGAGGGGCTTCAT-3’; mouse 18S: Forward: 5’ATTGGAGCTGGAATTACCGC-3’, Reverse: 5’CGGCTACCACATCCAAGGAA-3’.

### In vivo enteral treatment with CA7S

13-week-old male C57Bl/6J mice were purchased, acclimated, and housed as above. They were weight matched into two groups (p=0.88). After an overnight fast (17:00 to 0800), mice received either CA7S or PBS via direct duodenal and rectal administration. The optimal, physiologic dose of CA7S was extrapolated from the average pmol concentration of CA7S found in cecal samples from SG animals (average of 3000 pmol/mg of stool with 500 mg of stool per animal corresponds to 0.75 mg of CA7S per cecum).

Under isoflurane general anesthesia, 0.25 mg and 0.75 mg of CA7S in PBS (pH 7.2) was delivered by slow infusion (5 min) antegrade into the duodenum and retrograde into the rectum, respectively. The total volume of instillation was 2 mL (0.5 mg CA7S/mL). Control animals received similar volumes of PBS alone. 15 min post infusion, serum glucose was measured via tail vein followed by whole blood collection via cardiac puncture into K+EDTA tubes containing DPPIV inhibitor (Merck Millipore, Billerica, MA), Perfabloc (Sigma), and apoprotinin (Sigma). Organs were harvested for analysis. In order to account for changes in fasting times and hormonal diurnal rhythms, this experiment was carried out on four consecutive days such that only four mice were tested per day.

### In vivo CA7S gavage

16-week-old DIO mice were purchased and housed as described above. Mice were gavaged orally with 100 mg/kg CA7S from 20 mg/mL solution, or equivalent volume of PBS using 20G × 38mm gavage needle. 5 hours after CA7S/PBS administration, whole blood and intestinal segments were collected.

### In vivo CA7S and OGTT

Age matched, DIO mice were kept on HFD and their blood glucose levels were monitored until average fasting glucose levels were >160 mg/dL. Animals were fasted for 4 hours on the day of the experiment. Mice were matched into two groups based on fasting glucose levels and received either 100 mg/kg CA7S from a 20 mg/ml solution or an equivalent volume of PBS by oral gavage. Three hours later, an OGTT was performed using an oral gavage of 2 mg/g oral D-glucose (Sigma-Aldrich, St. Louis, MO). Blood glucose levels were measured at baseline and at minutes 15, 30, 60 and 120 with a OneTouch glucometer.

### Lentiviral IP injection

GLP-1R shRNA-containing lentiviral particles (LVP) were purchased from the MISSION TRC library (Sigma-Aldrich, St. Louis, MO). LVPs containing a mixture of three GLP-1R shRNA plasmid clones (TRCN0000004629, TRCN0000004630, and TRCN0000004633) were purchased, stored at −80 °C, and thawed on ice before use. DIO mice were maintained on a HFD until average fasting glucose >160 mg/dL in a BL2 facility. Under sterile conditions, mice were injected intraperitoneally with 0.2 ml of 5×10^5^ GLP-1R shRNA LVPs with a 27G needle^24,25^. 72 hours after LVP injection, mice underwent CA7S/PBS gavage followed by OGTT as above. After the OGTT was completed, mice were sacrificed and their tissues were harvested. GLP-1R knock-down efficiency was measured in tissues by qPCR as described above.

### Human stool collection

After obtaining institutional review board approval, we prospectively collected stool specimen from obese human subjects undergoing SG. Pre-operative stool specimen were collected on the day of surgery and post-operative stool specimen were obtained from post-operative day 14 to 99 (mode 15 days; median 36 days). Specimens were snap frozen in liquid nitrogen and stored at −80 °C until bile acid analysis was performed (as above).

## Acknowledgements

We thank members of the Devlin, Sheu, Clardy, and Banks labs for helpful discussions and advice. We would like to acknowledge the Blacklow and Kruse labs for help with equipment and reagents, and the BWH mouse facility. We are grateful to the human patients who participated in this study. This work was supported by a KL2 award from Harvard Catalyst (4Kl2TR001100-04) (E.G.S.), a pilot grant from Boston Area Diabetes and Endocrinology Research Center (BADERC) (NIH/NIDDK P30 DK057521) (E.G.S.), an NIH MIRA grant (R35 GM128618) (A.S.D.), a Blavatnik Biomedical Accelerator at Harvard University grant (A.S.D.), an American Heart Association Postdoctoral Fellowship (S.N.C.), an American College of Surgeons fellowship (D.A.H.), and an NIH T32 training grant (D.A.H.).

## Author contributions

A.S.D., E.G.S., S.N.C., and D.A.H. conceived the project and designed the experiments. S.N.C. performed the cell culture experiments, bile acid profiling, and transcriptional analyses and hormone quantifications on mouse tissues and blood. D.A.H. performed the surgeries and the enteral administration in vivo experiments. H.A. and R.S. performed the gavages, OGTTs, and the lentiviral injection experiments. M.T.H. performed NMR analyses. A.H.V. collected and provided the human samples. S.N.C., D.A.H., E.G.S., and A.S.D. wrote the manuscript. A.T. provided feedback and reviewed the manuscript. All authors edited and contributed to the critical review of the manuscript.

## Competing interests

CA7S is a subject of provisional patents held by HMS and BWH on which S.N.C., D.A.H., E.G.S., and A.S.D. are inventors. A.S.D. is a consultant for Kintai Therapeutics and HP Hood. E.G.S. was previously on the scientific advisory board of Kitotech, Inc.

## Data availability statement

All data generated or analyzed during this study are included in this article and its supplementary information files. In addition, numerical values for levels of individual bile acids shown as individual data points in Figure 2 and Supplementary Figures 3, 4, and 5 are available from the corresponding author upon reasonable request.

## Code availability statement

No custom code or mathematical algorithms were used in this study.

---

**Supplementary Figure 1.**
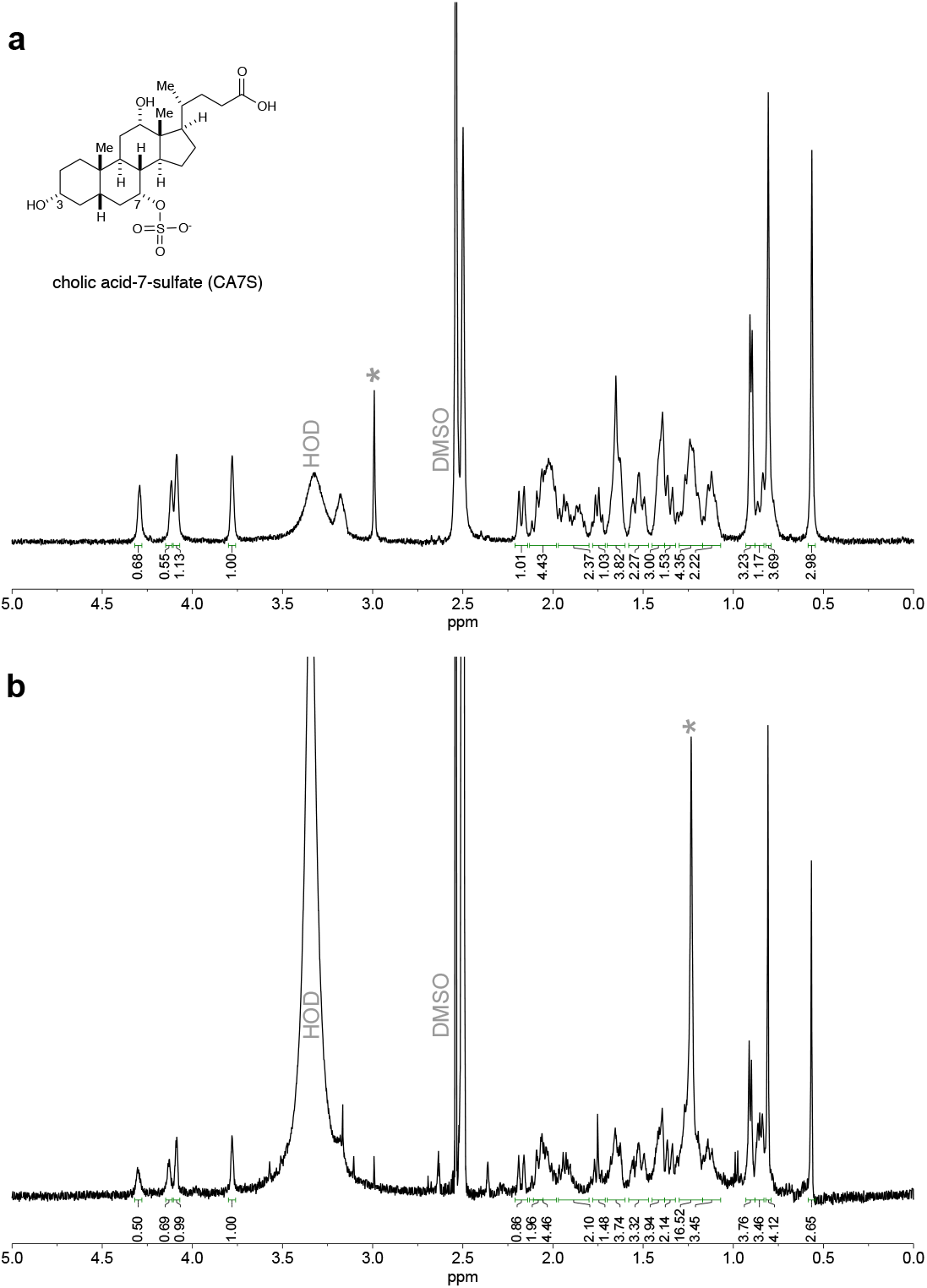
NMR of cholic acid-7-sulfate. **a**, ^1^H NMR of authentic sample of cholic acid-7-sulfate (Cayman Chemical). **b**, ^1^H NMR of CA7S purified from the cecal contents of SG mice. Signals between 3.7 to 4.4 ppm are diagnostic of CA7S. Impurities are denoted by asterisks.

**Supplementary Figure 2.**
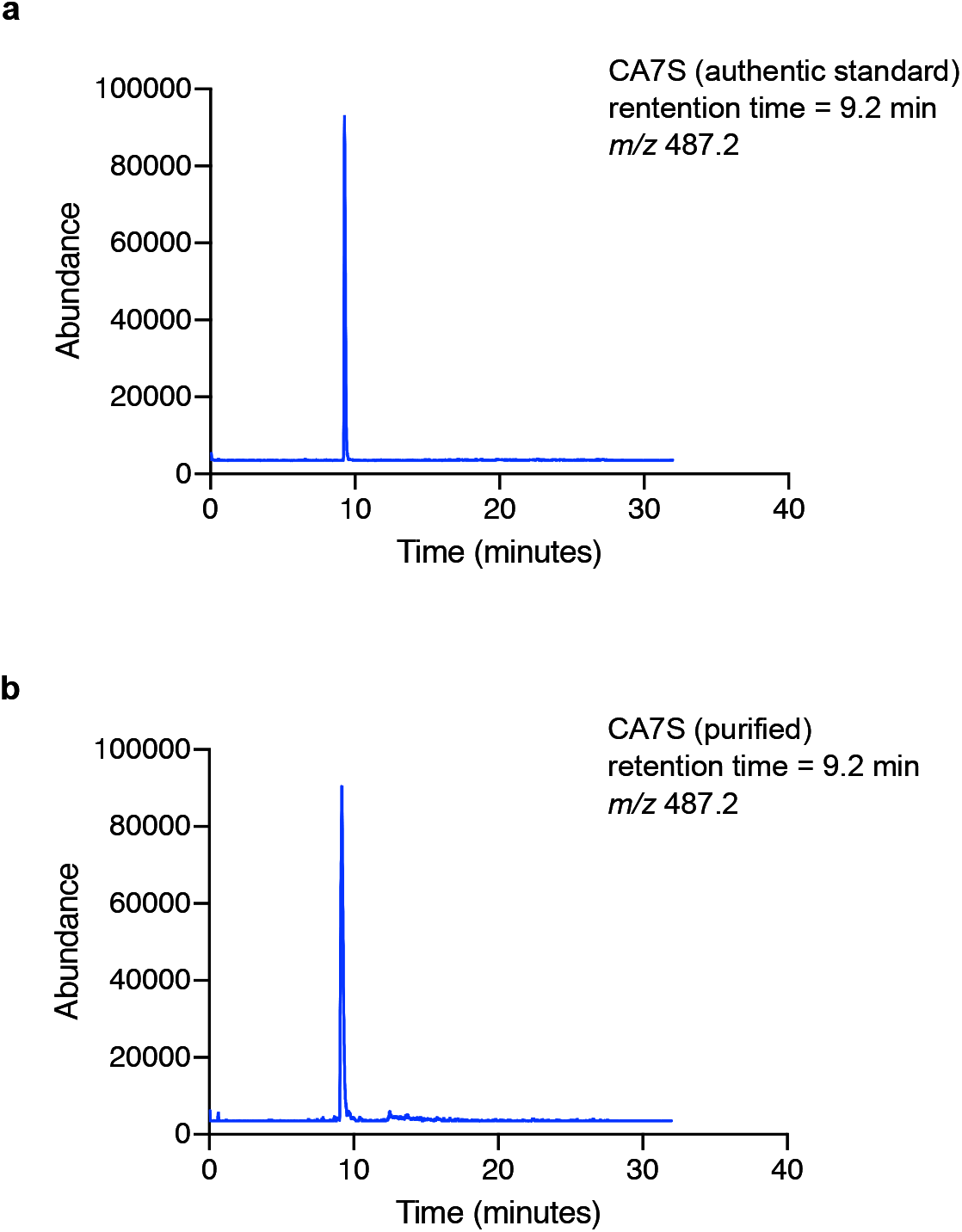
UPLC-MS analysis of cholic acid-7-sulfate. **a**, Commercially available cholic acid-7-sulfate (Cayman Chemical) and **b**, CA7S purified from the cecal contents of SG mice have the same mass (*m/z* 487.2) and elute at 9.2 minutes.

**Supplementary Figure 3.**
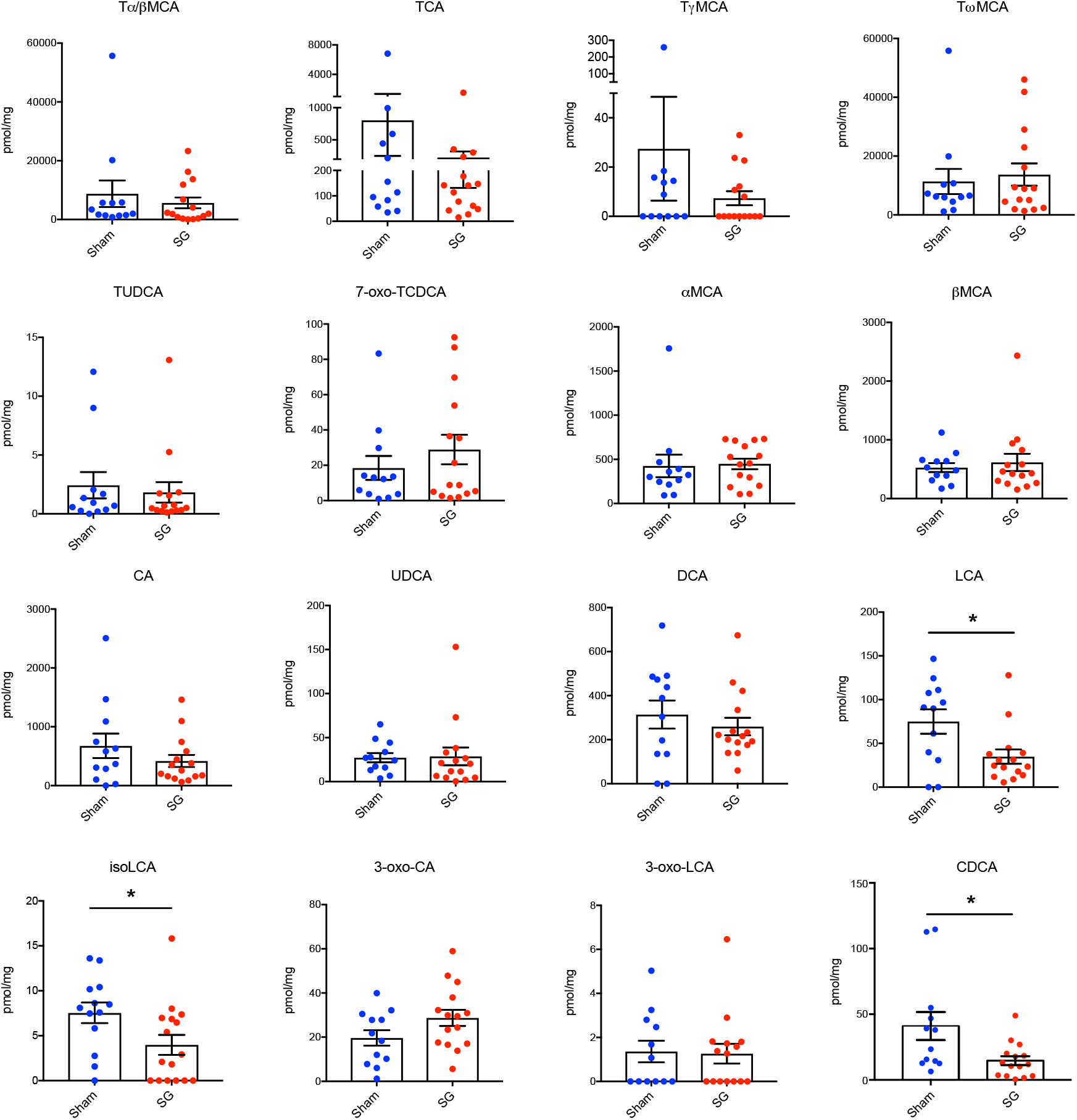
Bile acid concentrations in cecal contents of mice post-sham or post-SG. Cecal contents were collected from sham or SG mice 6 weeks post-op and bile acids were quantified using UPLC-MS (sham, n=12, SG, n=15, data not marked with asterisk(s) are not significant). All bile acids with measurable concentrations above the limit of detection are shown. Tα/βMCA, tauro-alpha- and tauro-beta-muricholic acid, *p*=0.537; TCA, tauro-cholic acid, *p*=0.325; TγMCA, tauro-gamma-muricholic acid, *p*=0.365; TωMCA, tauro-omega-muricholic acid, *p*=0.685; TUDCA, tauro-ursodeoxycholic acid, *p*=0.672; 7-oxo-TCDCA, 7-oxo-tauro-chenodeoxycholic acid *p*=0.34; αMCA, alpha-muricholic acid, *p*=0.874; βMCA, beta-muricholic acid, *p*=0.595; CA, cholic acid, *p*=0.287; UDCA, ursodeoxycholic acid, *p*=0.854; DCA, deoxycholic acid, *p*=0.481; LCA, lithocholic acid, \**p*=0.023; isoLCA, isolithocholic acid \**p*=0.024; 3-oxo-CA, 3-oxo-cholic acid, *p*=0.087; 3-oxo-LCA, 3-oxo-lithocholic acid, *p*=0.793; CDCA, chenodeoxycholic acid, \**p*=0.036, Welch’s t test. All data are presented as mean ± SEM.

**Supplementary Figure 4.**
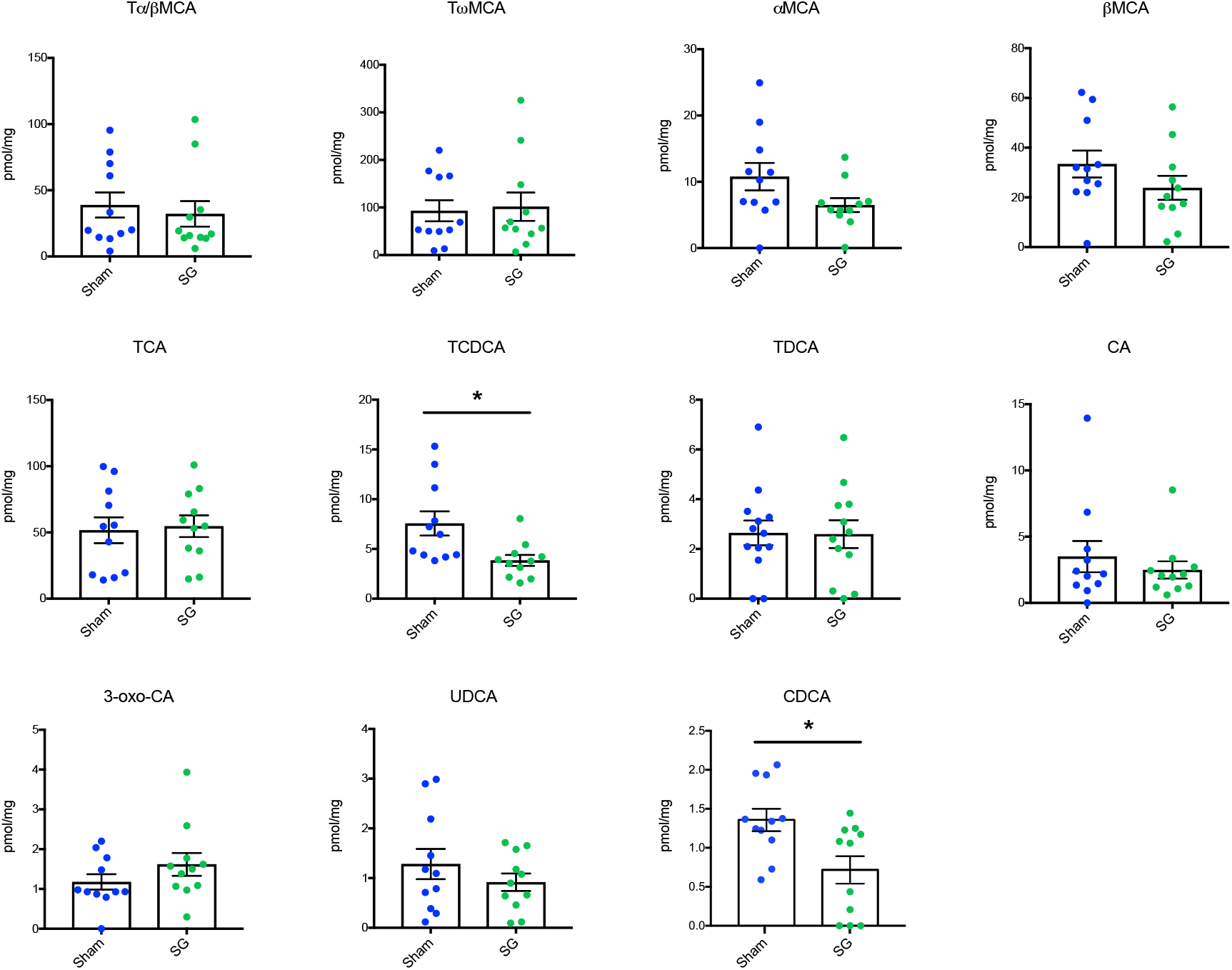
Bile acid concentrations in livers of mice post-sham or post-SG. Livers were collected from sham or SG mice 6 weeks post-op and bile acids were quantified using UPLC-MS (n=11 per group, data not marked with asterisk(s) are not significant). All bile acids with measurable concentrations above the limit of detection are shown. Tα/βMCA, tauro-alpha- and tauro-beta-muricholic acid, *p*=0.625; TωMCA, tauro-omega-muricholic acid, *p*=0.822; αMCA, alpha-muricholic acid, *p*=0.085; βMCA, beta-muricholic acid, *p*=0.203; TCA, tauro-cholic acid, *p*=0.815; TCDCA, tauro-chenodeoxycholic acid, \**p*=0.015; TDCA, tauro-deoxycholic acid, *p*=0.947; CA, cholic acid, *p*=0.468; 3-oxo-CA, 3-oxo-cholic acid, *p*=0.22; UDCA, ursodeoxycholic acid, *p*=0.314; CDCA, chenodeoxycholic acid, \**p*=0.023, Welch’s t test. All data are presented as mean ± SEM.

**Supplementary Figure 5.**
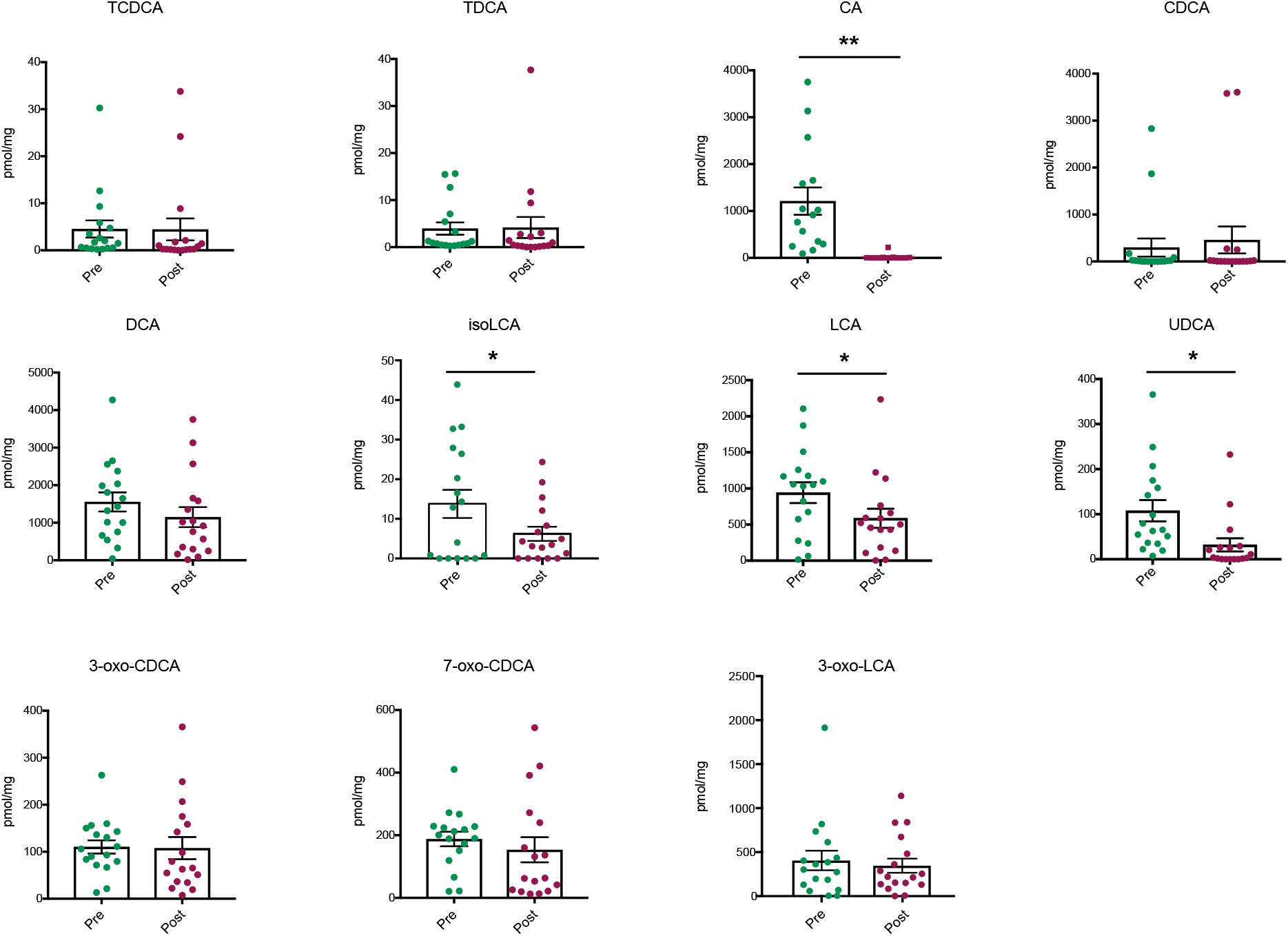
Bile acid concentrations in feces of human patients pre-SG or post-SG. Feces were collected from patients pre-op or ~5 weeks post-op and bile acids were quantified using UPLC-MS (n=17 patients, median 36 days after surgery, data not marked with asterisk(s) are not significant). All bile acids with measurable concentrations above the limit of detection are shown. TCDCA, tauro-chenodeoxycholic acid, *p*=0.973; TDCA, tauro-deoxycholic acid, *p*=0.933; CA, cholic acid, \*\**p*=0.001; CDCA, chenodeoxycholic acid, *p*=0.525; DCA, deoxycholic acid, *p*=0.135; LCA, lithocholic acid, \**p*=0.014; isoLCA, iso-lithocholic acid, \**p*=0.032; UDCA, ursodeoxycholic acid, \**p*=0.022; 3-oxo-CDCA, 3-oxo-chenodeoxycholic acid, *p*=0.921; 7-oxo-CDCA, 7-oxo-chenodeoxycholic acid, *p*=0.477, 3-oxo-LCA, 3-oxo-lithocholic acid, *p*=0.563, paired t test. All data are presented as mean ± SEM.

**Supplementary Figure 6.**
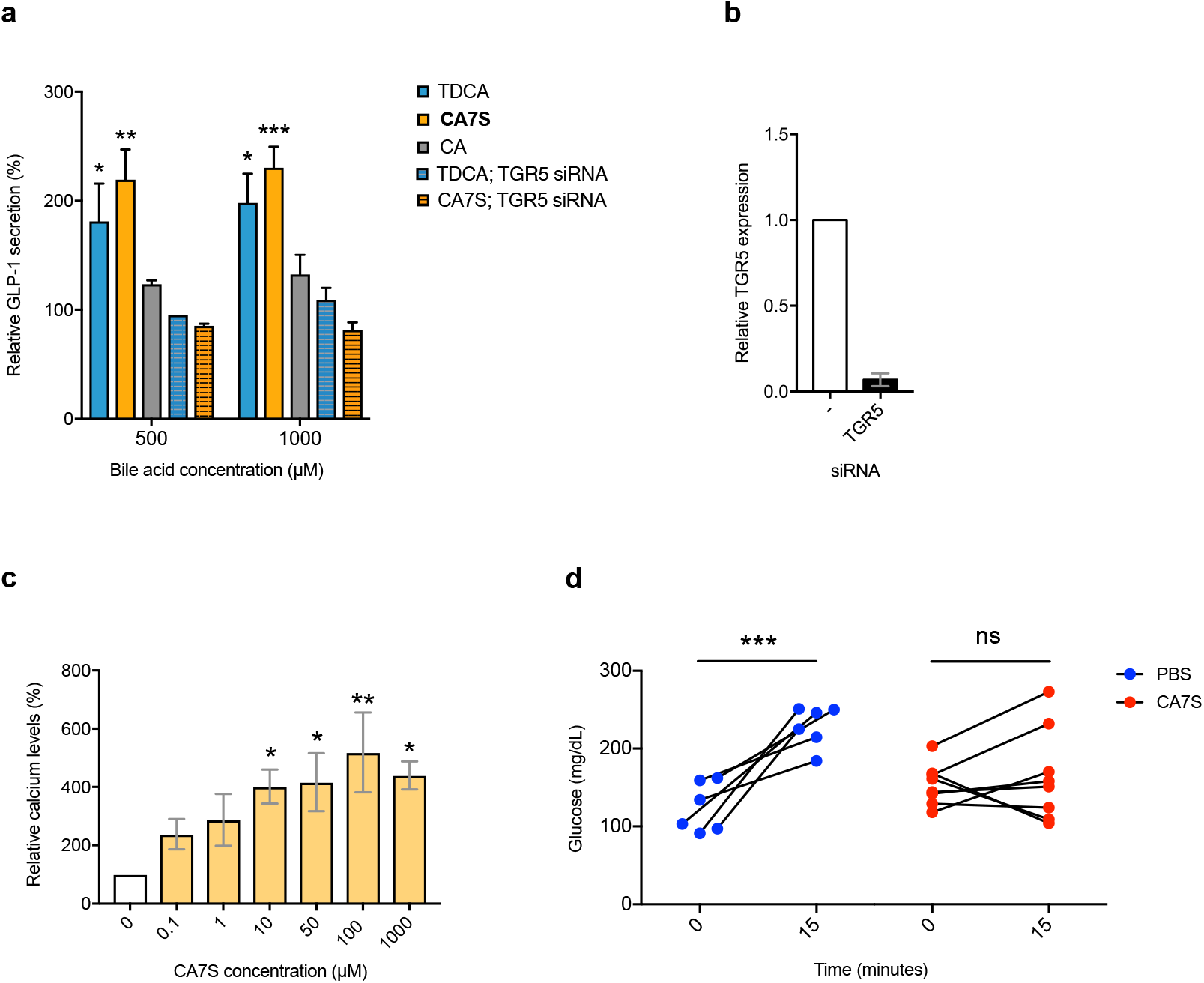
CA7S activates TGR5, induces GLP-1 secretion, and reduces systemic glucose levels. **a**, CA7S induced secretion of GLP-1 in NCI-H716 cells compared to both CA and the known TGR5 agonist, TDCA. siRNA-mediated knockdown of TGR5 abolished GLP-1 secretion (≥3 biological replicates per condition, \**p*=0.027-0.04, \*\**p*=0.003, \*\*\**p*=0.0007, one-way ANOVA followed by Dunnett’s multiple comparisons test). **b**, Quantitative real time PCR analysis of expression of human TGR5 in TGR5 siRNA and negative (−) siRNA-treated NCI-H716 cells for Fig. 2g and Supplementary Fig. 6a. **c**, CA7S induced an increase in intracellular calcium levels in NCI-H716 cells (≥3 biological replicates per condition, \**p*=0.018-0.041, \*\**p*=0.003, one-way ANOVA followed by Dunnett’s multiple comparisons test). **d**, In vivo change in serum glucose upon treatment with PBS and CA7S (PBS, n=6; CA7S, n=8 mice per group, \*\*\**p*=0.0001, ns=not significant *p*=0.63, paired t test). All data are presented as mean ± SEM.

**Supplementary Figure 7.**
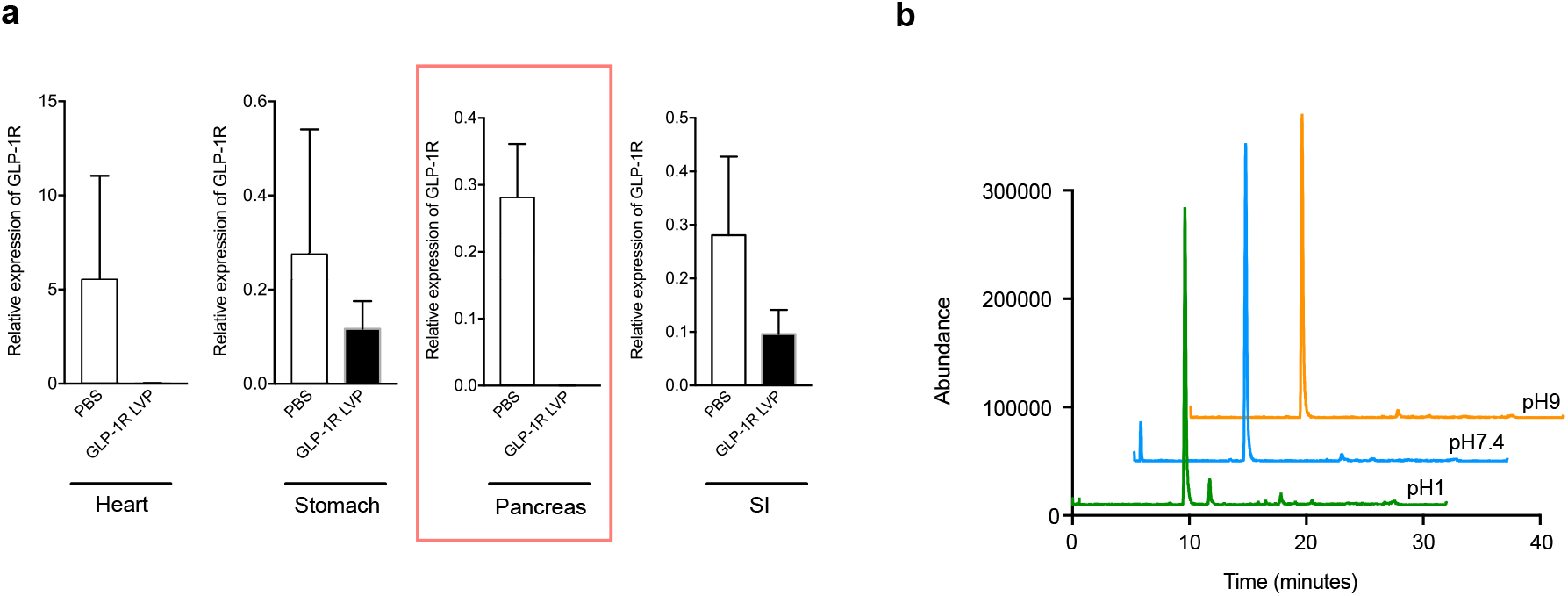
GLP-1R shRNA knockdown efficiency and stability of CA7S. **a**, Quantitative real time PCR analysis in mice corresponding to Fig. 4f and 4g. Animals were injected with lentiviral shRNA targeting GLP-1R or PBS. Expression of mouse GLP-1R in indicated tissues of mice was measured following OGTT, which was performed 3 days post-injection (SI = small intestine, PBS, n=2; GLP-1R LVP shRNA n=14). **b**, UPLC-MS traces of CA7S after incubation at 37 °C in buffer at the indicated physiological pHs. All data are presented as mean ± SEM.

